# Disentangling primer interactions improves SARS-CoV-2 genome sequencing by the ARTIC Network’s multiplex PCR

**DOI:** 10.1101/2020.03.10.985150

**Authors:** Kentaro Itokawa, Tsuyoshi Sekizuka, Masanori Hashino, Rina Tanaka, Makoto Kuroda

## Abstract

Since December 2019, the coronavirus disease 2019 (COVID-19) caused by a novel coronavirus SARS-CoV-2 has rapidly spread to almost every nation in the world. Soon after the pandemic was recognized by epidemiologists, a group of biologists comprising the ARTIC Network, has devised a multiplexed polymerase chain reaction (PCR) protocol and primer set for targeted whole-genome amplification of SARS-CoV-2. The ARTIC primer set amplifies 98 amplicons, which are separated only in two PCRs, across a nearly entire viral genome. The original primer set and protocol showed a fairly small amplification bias when clinical samples with relatively high viral loads were used. However, when sample’s viral load was low, several amplicons, especially amplicons 18 and 76, exhibited low coverage or complete dropout. We have determined that these dropouts were due to a dimer formation between the forward primer for amplicon 18, 18_LEFT, and the reverse primer for amplicon 76, 76_RIGHT. Replacement of 76_RIGHT with an alternatively designed primer was sufficient to produce a drastic improvement in coverage of both amplicons. Based on this result, we replaced 12 primers in total in the ARTIC primer set that were predicted to be involved in 14 primer interactions. The resulting primer set, version N1 (NIID-1), exhibits improved overall coverage compared to the ARTIC Network’s original (V1) and modified (V3) primer set.

## Background

The realtime surveillance of pathogen genome sequences during an outbreak enables monitoring of numerous epidemical factors such as pathogen adaptation and transmission chains in local to even global scale (Gardy and Loman 2017, Hadfield et al. 2018). Since it was first identified in Hubei, China in December 2019 (Zhu et al. 2020), the novel coronavirus, SARS-CoV-2, responsible for the atypical respiratory illness COVID-19, has become a major concern for the medical community around the world. The relatively large genome size of corona viruses (approx. 30 kb) and varying levels of viral load in clinical specimens have made it challenging to reconstruct the entire viral genome in a simple and cost-effective manner. In January 2020, a group of biologists comprising the ARTIC Network (https://artic.network/), designed 196 primer (98 pairs) (https://github.com/artic-network/artic-ncov2019/tree/master/primer_schemes/nCoV-2019/V1) for targeted amplification of the SARS-CoV-2 genome by multiplexing PCR. These primers and method were based on a primer design tool Primal Scheme and a laboratory protocol PrimalSeq that had been previously developed for sequencing outbreaking RNA virus genomes directly from clinical samples using portable nanopore sequencer or other NGS platforms (Quick et al. 2017, Grubaugh et al. 2019). The ARTIC primer set for SARS-coV-2 (hereafter, ARTIC primer set V1) is designed to tile amplicons across nearly entire sequence of the published reference SARS-CoV-2 genome MN908947.3 (Wu et al. 2020). The 98 primer pairs are divided into two separate subsets (Pools 1 and 2), such that no overlap between PCR fragments occurs in the same reaction.

The ARTIC primer set V1 and the published protocol (Quick 2020) worked quite for samples with a relatively high viral load (Ct < 25 in clinical qPCR tests). For these samples, all designated amplicons are amplified with an acceptable level of coverage bias for subsequent NGS analysis. However, a gradual increase in the overall coverage bias was observed as a sample’sviral load decreased. Although this phenomenon is generally expected in such highly multiplexed PCR, the coverage for the two particular PCR amplicons, 18 and 76, which correspond to regions of genome encoding nsp3 in ORF1a and the spike (S) protein, respectively, decays far more rapidly than other targets (Fig 1A). In our experience, the low to zero depth for those two amplicons was the most frequent bottleneck for using the ARTIC primer set V1 to sequence all targeted genomic regions from samples with middle to low viral load (Ct > 27). In situation with a high coverage bias in genome sequencing such as seen in Fig 1A, excessive sequencing efforts are required to obtain viral genome sequences with no or few gaps. Thus, minimizing the overall coverage bias will benefit the research community by both enabling more multiplexing in given sequencing capacity and lowering the sequencing cost per sample.

**Fig 1.**
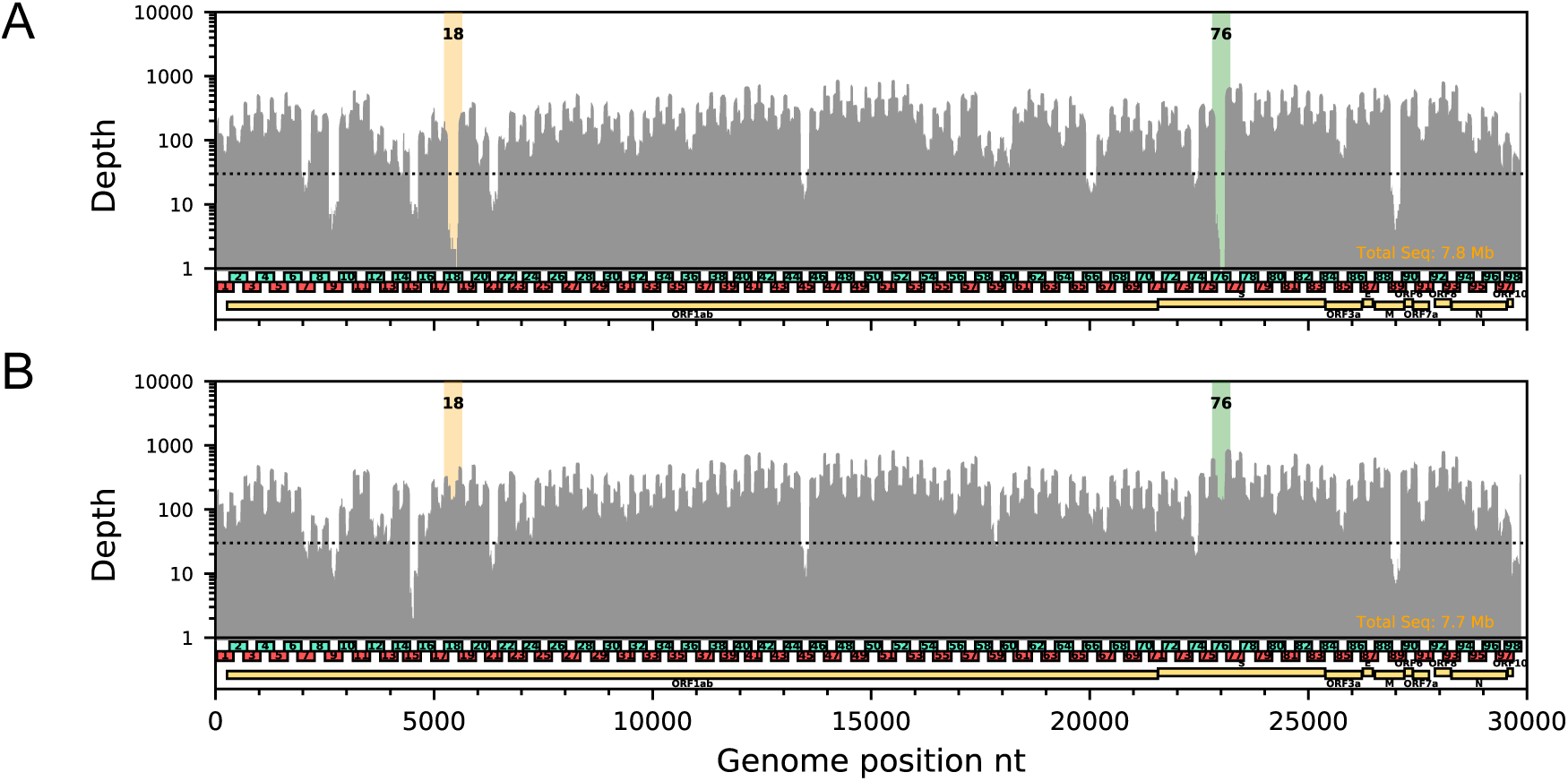
Examples of depth plot for (A) original ARTIC primer set V1 and (B) V1 with 76_RIGHT replacement for the same clinical sample (previously deposited to GISAID with ID EPI_ISL_416596, Ct=28.5, 1/25 input per reaction). Regions covered by amplicons with modified primer (76_RIGHT) and the interacting primer (18_LEFT) are highlighted by green and orange colors, respectively. For all data, reads were downsampled to normalize average coverage to 250X. Horizontal dotted line indicates depth = 30. These two experiments were conducted with the same PCR master mix (except primers) and in the same PCR run in the same thermal cycler.

In this report, we first show that the acute dropout of amplicons 18 and 76 was due to the formation of a single dimer between the forward primer for amplicon 18 and the reverse primer for amplicon 76. The replacement of one of the two interacting primers resolved the dropout of both amplicons. We further detected an additional 13 other potential primer interactions that may be responsible for low coverage in other regions covered by the affected amplicons. Our modified primer set, version N1 (NIID-1), which includes 12 primer replacements from the ARTIC primer set V1, yielded improved overall genome coverage in clinical samples compared to V1 and another modified primer set, V3. The results indicated that preventing primer dimer-formation is an effective measure to improve coverage bias in the ARTIC Network’s SARS-CoV-2 genome sequencing protocol, and may be applicable to other PrimalSeq methods in general.

## Results and Discussion

In the original ARTIC prime set V1, PCR amplicons 18 and 76 were amplified by the primer pairs 18_LEFT & 18_RIGHT and 76_LEFT & 76_RIGHT, respectively. Those primers were included in the same multiplexed reaction, “Pool 2.” We noticed that two of those primers, 18_LEFT and 76_RIGHT, were perfectly complementary to one another by 10-nt at their 3′ ends (Fig 1). Indeed, we observed NGS reads derived from the predicted dimer in raw FASTQ data. From this observation, we reasoned that the acute dropouts of those amplicons were due to an interaction between 18_LEFT and 76_RIGHT, which could compete for amplification of the designated targets. Next, we replaced one of the two interacting primers, 76_RIGHT, in the Pool 2 reaction with a newly designed primer 76_RIGHTv2 (5′-TCTCTGCCAAATTGTTGGAAAGGCA-3′), which is located 48-nt downstream from 76_RIGHT. Figures 1A and 1B show the coverage obtained with the V1 set and the V1 set with 76_RIGHT replaced with 76_RIGHTv2 for cDNA isolated from a clinical sample obtained during the COVID-19 cruise ship outbreak, which was previously analyzed (EPI_ISL_416596) (Sekizuka et al. 2020). The replacement of the primer drastically improved the read depth in the regions covered by amplicons 18 and 76 without any notable adverse effects. The replacement of the primer 76_RIGTH improved coverage not only for amplicon 76, but also for 18 as well, supporting the hypothesis that the single primer interaction caused dropout of both amplicons.

Given this observation, we identified an additional 13 primer interactions using *in silico* analysis (Fig 2A and B). Those primer interactions predicted by PrimerROC algorithm (Johnston et al. 2019), which gave the highest score for the interaction between 18_LEFT and 76_RIGHT among all 4,743 possible interactions, were likely involved in producing the low coverage frequently seen in our routine experiments. Next, we designed an additional 11 alternative primers, which resulted in a new primer set (ARTIC primer set ver. NIID-1 (N1) including 12 primer replacements from the original V1 primer set (Table S1). The N1 primer set eliminated all interactions shown in Fig 2A, and was expected to improve amplification of up to 22 amplicons (1, 7, 9, 13, 15, 18, 21, 29, 31, 32, 36, 38, 45, 48, 54, 59, 66, 70, 73, 76, 85, and 89). Alongside with this modification, the ARTIC Network itself released another modified version of primer set known as V3 in 24^th^ March 2020 (Loman and Quick 2020) after we reported our result on the replacement of primer 76_RIGHT in a preprint(Itokawa et al. 2020a). The V3 primer set included 22 spike-in primers, which were directly added into the V1 primer set to aid amplification of 11 amplicons (7, 9, 14, 15, 18, 21, 44, 45, 46, 76, and 89).

**Fig 2.**
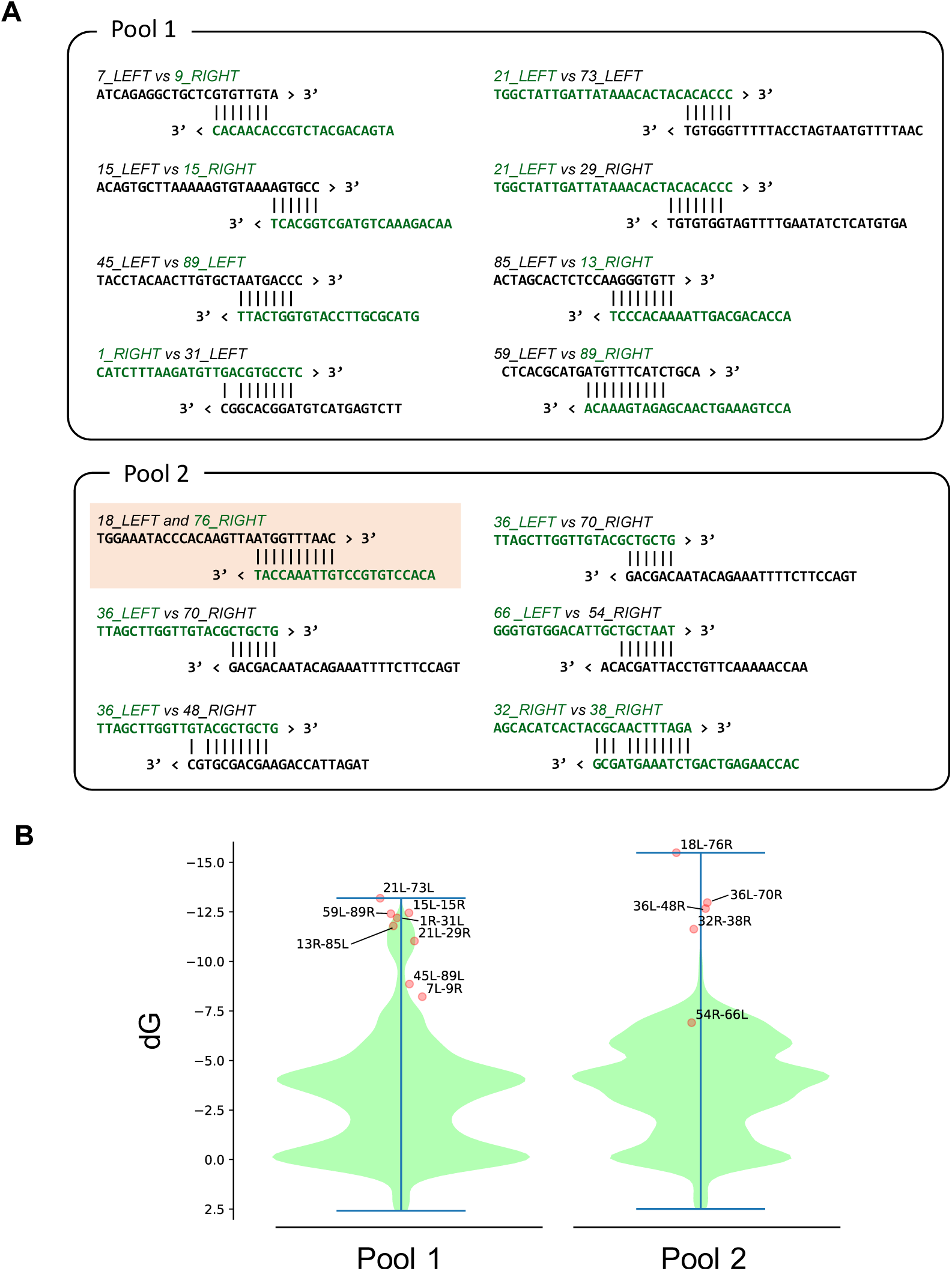
(A) The 14 predicted primer interactions subjected for modification in this study. Primers replaced in the N1 primer set (Table 1) are shown in green. (B) Violin plots showing the distributions of dimer scores (dG) at all heterodimers in pools1 and 2 reported by the PrimerROC algorithm (n=4,743 for each pool). The scores of interactions depicted in Fig 2A are over plotted as scatter points.

We compared the performance of the original primer set (V1) and the two modified primer sets (V3 and N1) by observing their responses to different annealing/extension temperatures () in the thermal program (98 °C for 30 s followed by 30 cycles of 98 °C for 15 s and *Ta* °C for 5 min) using the gradient function of a thermal cycler. We surmised that this gradient temperature experiment would enable us to examine the dynamics of amplification efficiencies for each amplicon over varying annealing condition. In general, amplicons suffering from primer interactions were expected to drop rapidly as *Ta* decreases. Figure 3 indicates the abundances of the 98 amplicons at eight different *Ta*, ranging from 63.1–68.6 °C, using same dilution from a cDNA sample with high viral load (Ct = 16.0), which has previously been obtained from patients during the cruise ship outbreak sequenced (EPI_ISL_416584). With the V1 primer set, amplicons 18 and 76 exhibited extremely low coverage for all *Ta* values, with only a slight improvement above 67 °C. In addition to those two amplicons, many other amplicons exhibited reduced coverage in thee lower *Ta* range. Most of those amplicons were related to the predicted primer interactions depicted in figure 2A. Although the dropout for amplicons 18 and 76 resolved with the V3 primer set, many amplicons still suffered low coverage in the low *Ta* region. Compared to the V1 and V3 primer sets, the modifications in the N1 primer set resulted in improved robustness of coverage over a broader *Ta* range for relevant amplicons. The improvement, however, made potentially weak amplicons 74 and 98 more apparent (Fig 3). The abundance of amplicons 74 gradually decreased with decreasing *Ta*, in contrast, the abundance of amplicon decreased with increasing *Ta*. These amplicons seemed equally weak in all three primer sets rather than specific in N1 primer set. So far, we have not yet identified interactions involving the primers for those amplicons. The gradient experiment also revealed relatively narrow range of optimal temperature for *Ta* for the V1 and V3 primer set, around 65 °C, which was broaden for the N1 primer set. Nevertheless, while Ta = 65 °C is a good starting point, a fine tuning of this value may help improving sequencing quality since even slight difference between thermal cyclers, such as systematic and/or well-to-well accuracy differences and under-or overshooting, may affect the results of multiplex PCR (Ho Kim et al. 2008). Finally, we further compared the V1, V3 and N1 primer sets for three other clinical samples using a standard temperature program (*Ta* = 65 °C). In all three clinical samples (Fig 4 and S1), the N1 primer set showed the most even coverage distribution.

**Fig 3.**
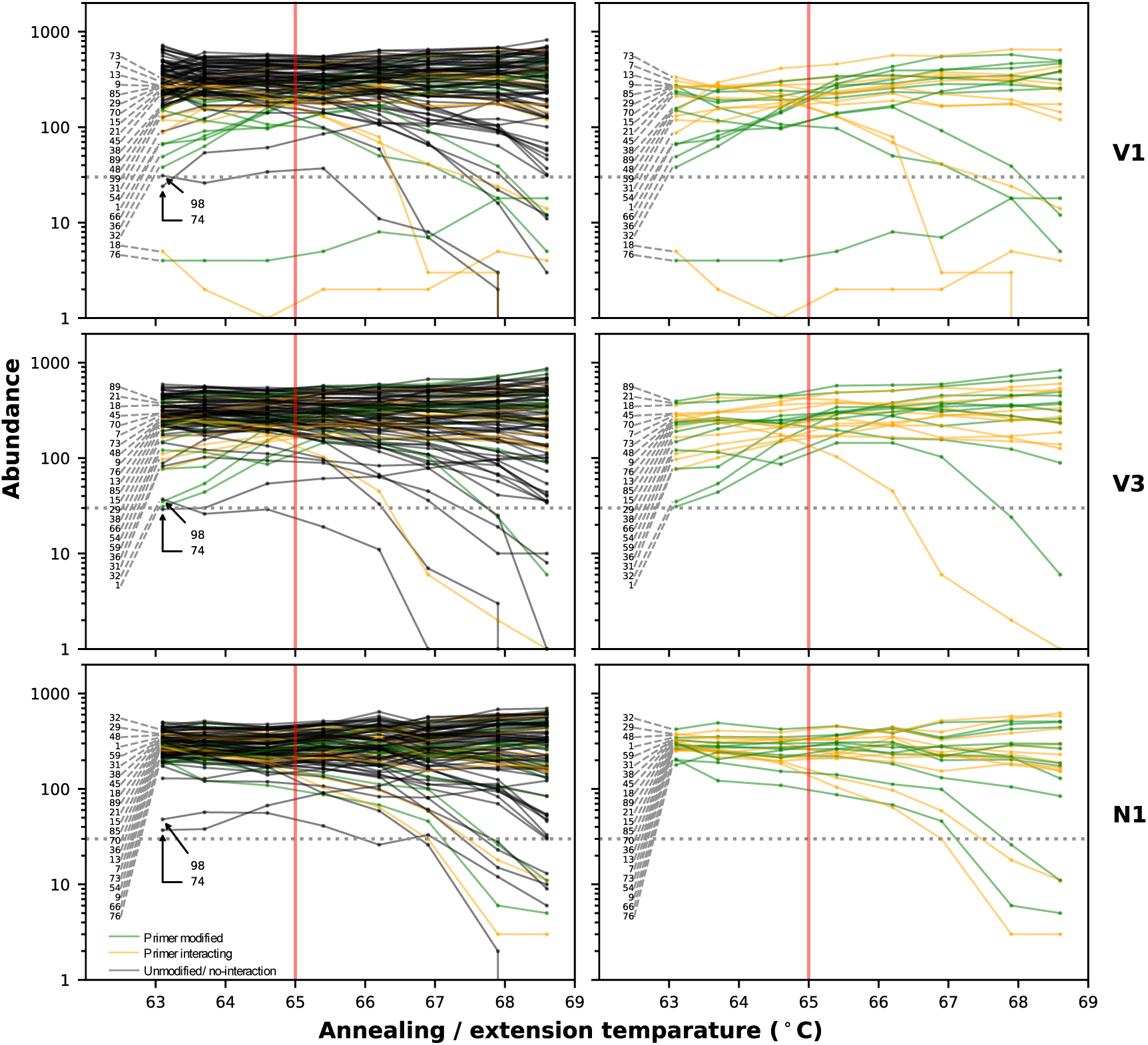
Abundance of 98 amplicons at 8 different annealing/extension temperatures with the three different primer sets on a same clinical sample (previously deposited to GISAID with ID EPI_ISL_416584, Ct=16, 1/300 input per reaction). For all data, reads were downsampled to normalize average coverage to 500X before analysis. The green lines and points indicate the abundances of amplicons whose primers in V1 primer set were subjected to modification in the N1 primer set. The orange lines and points indicate the abundances of amplicons whose primers were not modified but predicted to be eliminated the adverse primer interactions in the N1 primer set. Other amplicons which were not subjected to the modification are indicated by black lines and points. The plots in the left column shows results of all 98 amplicons while only amplicons targeted by modification are shown in the plots in the right column. Horizontal dotted line indicates fragment abundance = 30. Red vertical lines indicate normal annealing/extension temperature, 65 °C. All those experiments were conducted with the same PCR master mix (except primers) and in the same PCR run in the same thermal cycler.

**Fig 4.**
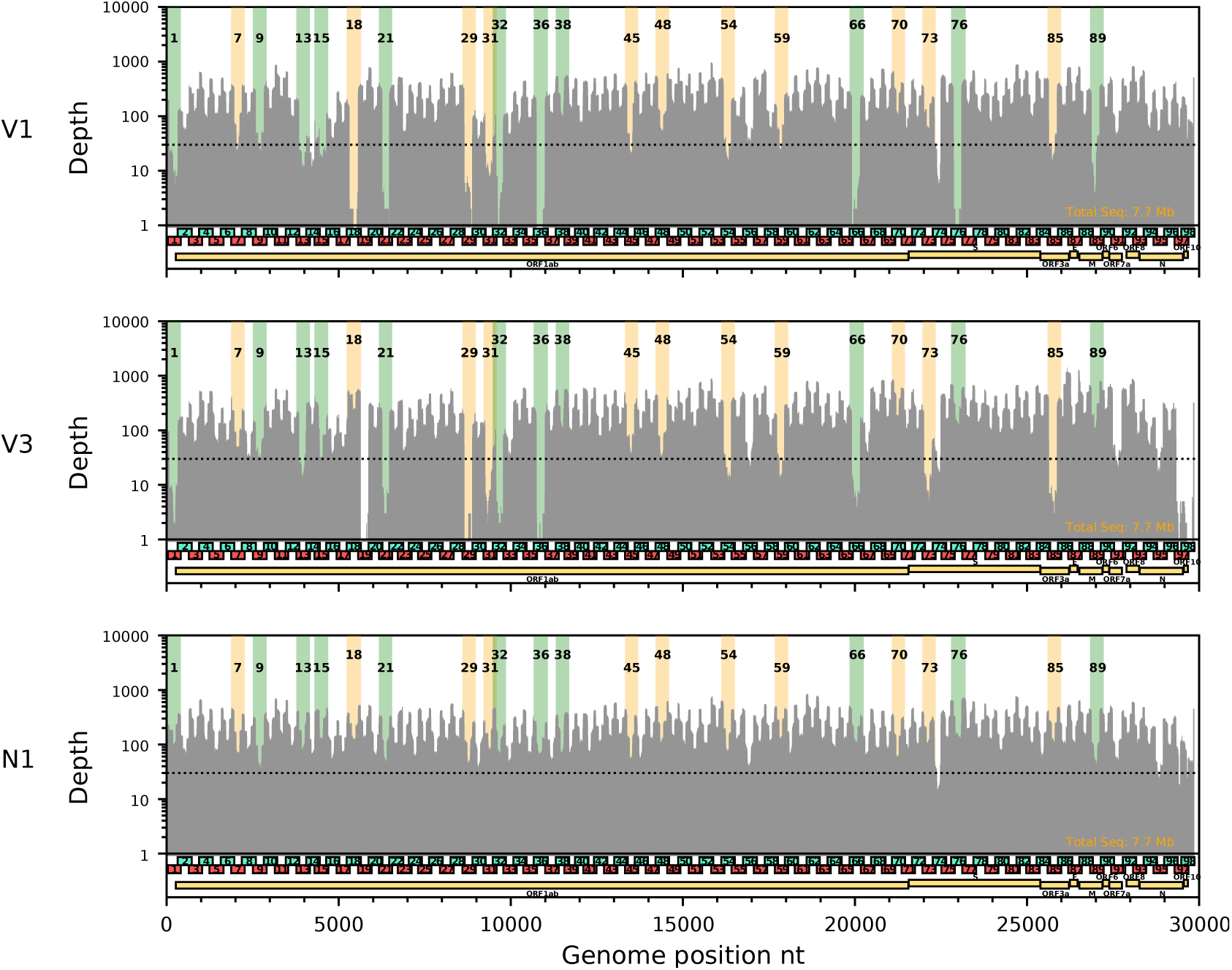
A depth plot of original (V1) and two modified ARTIC primer sets (V3 and N1) on a same clinical sample (previously deposited to GISAID with ID EPI_ISL_416596, Ct=28.5, 1/25 input per reaction). Regions covered by amplicons with modified primers and with not modified but interacting primers are highlighted by green and orange colors, respectively. For all data, the reads were downsampled to normalize average coverage to 250X. Horizontal dotted line indicates depth = 30. These two experiments were conducted with the same PCR master mix (except primers) and in the same PCR run in the same thermal cycler.

## Conclusions

The formation of primer-dimers is a major cause of coverage bias in the ARTIC Network’s multiplex PCR protocol for SARS-CoV-2 genome sequencing. Eliminating these problematic primer interactions improves sequence coverage and will likely increase the quality of genome sequencing.

## Materials and Methods

### Design of alternative primers

Re-design of the primer nCoV-2019_76_RIGHT was done using PRIMER3 software (Untergasser et al. 2012). Other primers were basically re-designed just by shifting their position several nucleotides toward the 5′ ends, but extension or trimming on either end were applied when the medium dissociation temperature (*Tm*) predicted by the NEB website tool (https://tmcalculator.neb.com/) were considered too low or high. See details of modifications on primers indicated in Table S1. All new primers were assessed by PrimerROC (Johnston et al. 2019) (http://www.primer-dimer.com/) to ensure no significant interactions with the remaining primers were predicted. All primer sequences included in the primer set N1 and information for their genomic positions were deposited to https://github.com/ItokawaK/Alt_nCov2019_primers. All primers used in this study were synthesized as OPC purification grade by Eurofins Genomics in Japan.

### cDNA samples and multiplex PCR

Four cDNA samples obtained from clinical specimens (pharyngeal swabs) during the COVID-19 outbreak on a cruise ship February 2020 (Sekizuka et al. 2020) were reused in this study. The cDNA had been synthesized by a reverse transcription protocol published by the ARTIC Network (Quick 2020) and diluted to 5-fold by H_2_O. For the temperature gradient experiment in figure 3, a cDNA from a very high viral load (Ct = 16.0) was further diluted 25-fold by H_2_O and used for the PCR reactions. This dilution and scaling down of the PCR reaction volume, as described below, made the amount of input cDNA approx. 1/300 per reaction compared to the original ARTIC Network’s protocol (Quick 2020). For experiments depicted in Fig 1 and 4, three cDNA from clinical samples with moderate viral loads (Ct = 26.5−28.5) was further diluted 2-fold with H_2_O to allow for multiple experiments on the same sample. This dilution and the scaling down of PCR reaction volume made the amount of input cDNA approx. 1/25 per reaction compared to the original ARTIC Network’s protocol. For the multiplex PCR reactions, 1 μl of the diluted cDNA was used in 10 μl reaction mixture of Q5 Hot START DNA Polymerase kit (NEB) (2 μl of 5X buffer, 0.8 μl of 2.5 mM dNTPs, 0.1 μl of polymerase and 0.29 μl of 50 μM primer mix, adjusted by milli-Q water to 10 μl). The thermal program was identical to the original ARTIC protocol: 30 sec polymerase activation at 98 °C followed by 30 cycles of 15 sec denaturing at 98 °C and 5 min annealing and extension at 65 °C (or variable values in gradient mode) in Thermal Cycler Dice ® (Takara Bio). The PCR products in Pool 1 and 2 reactions for same clinical samples were combined and purified with 1X concentration of AmpureXP.

### Sequencing

The purified PCR product was subjected to illumina library prep using QIAseq FX library kit (Qiagen) in 1/4 scale and using 6 min fragmentation time (Itokawa et al. 2020b). After the ligation of barcoded adaptor, libraries were heated to 65 °C for 20 min to inactivate the ligase, and then all libraries were pooled in a 1.5 ml tube. The pooled library was first purified by AmpureXP at 0.8X concentration, and then again at 1.2X concentration. The purified library was sequenced for 151 cycles at both paired-ends in Illumina iSeq100.

### Coverage and depth analysis

The obtained reads were mapped to the reference genome of SARS-CoV-2 MN908947.3 (Wu et al. 2020) using *bw mem* (Li and Durbin 2009). The *epth* function in *s mtools* (Li et al. 2009) with ‘aa’ option was used to determine coverage at each base position. Then, the ‘s’ option of *s mtools iew* function was used for subsampling reads from each mapping data for normalization.

The coverage (fragment abundance) analysis was conducted by defining 98 representative small regions (10-nt) that are unique to the 98 amplicons amplified by either the V1 or N1 primer set (Supportive file; amplicon_representative_regions.bed). Those representative small regions also reside within overlapped regions where original and corresponding alternative primers in the V3 primer set amplify (e.g. one region overlap with both amplicons amplified by 7_LEFT & 7_RIGHT and 7_LEFT_alt0 & 7_LEFT_alt5). The start and end mapping positions of whole insert sequences of paired-end reads were determined by the *b mtobe* function in bedtools (Quinlan and Hall 2010) with ‘bedpe’ option. The number of insert fragments overlapping each defined small representative region were counted by the *co er ge* function in the bedtools. Fragment inserts of unexpectedly long length (>500 bp from start to end positions) were filtered out from the analysis. The depth counts were summarized and visualized using the python3.6 and the *m tplotlib* library (Hunter 2007).

## Author contributions

KI designed the new primers and drafted this manuscript. KI and TS analyzed data. KI, MH and RT performed all experiments. MK and TK critically reviewed the manuscript.

## Ethics statement

This study was approved by the research ethics committee of the National Institute of Infectious Diseases (approval no. 1091). It was conducted according to the principles of the Declaration of Helsinki, in compliance with the Law Concerning the Prevention of Infections and Medical Care for Patients of Infections of Japan. The ethical committee waived the need for written consent regarding the research into the viral genome sequence. The personal data related to the clinical information were anonymized, and our procedure is not to request written consent for all patients suffering from COVID-19.

## Acknowledgments

First, we must note the huge contribution of the ARTIC Network, who developed the original primer set V1 and immediately opened it to the community. We deeply thank the staff of the Quarantine Station for the collection and transport of clinical specimens and the staff of the Influenza Virus Research Center and Department of Virology 3, National Institute of Infectious Diseases for providing clinical samples used in this study..

## Supportive files

**Table_S1.pdf** Differences between the N1 primer set and the original V1 primer set.

**Fig_S1.pdf** Depth plots of the original (V1) and two modified ARTIC primer sets (V3 and N1) for two clinical samples (newly deposited to GISAID with ID EPI_ISL_416749, Ct = 27.3 for A and previously deposited with ID EPI_ISL_416596, Ct = 26.5 for B, each 1/25 input per reaction). Regions covered by amplicons with modified primers and with not modified but interacting primers are highlighted by green and orange colors, respectively. For all data, reads were downsampled to normalize average coverage to 250X. Horizontal dotted line indicates depth = 30. These two experiments were conducted with the same PCR master mix (except primers) and in the same PCR run in the same thermal cycler.

**amplicon_representative_regions.bed** BED format file for representative small regions used for calculating amplicon abundances.

