## Supplementary figures and images for "Disentangling primer interactions improves SARS-CoV-2 genome sequencing by the ARTIC Network’s multiplex PCR"

### Figure S1

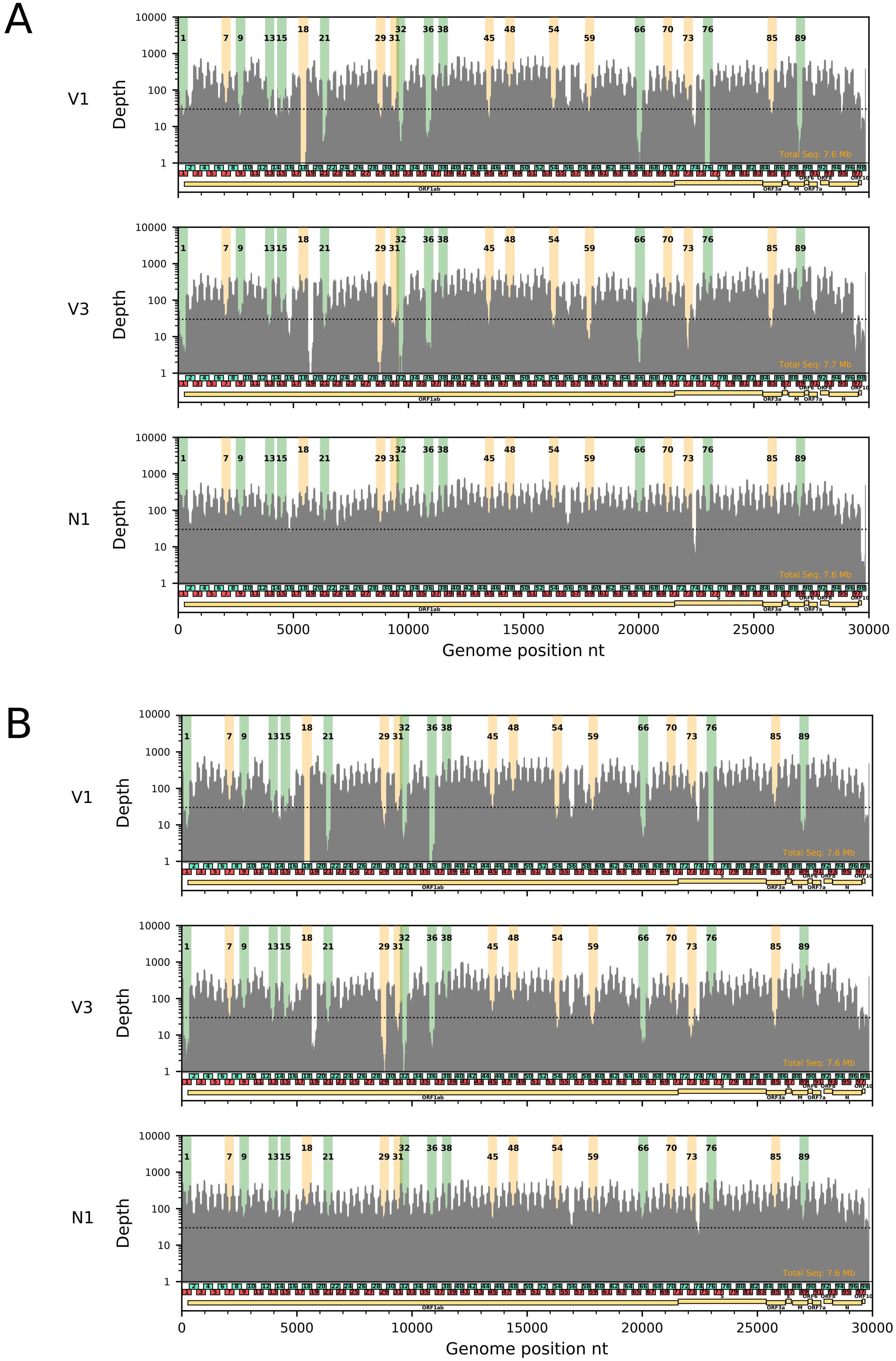
