## Supplementary material for "Disentangling primer interactions improves SARS-CoV-2 genome sequencing by the ARTIC Network’s multiplex PCR": Table S1

Table S1 Difference of the N1 primer set to the original V1 primer set

| Primer name (nCoV-2019_***) | Operation | Location (1-base) | Direction | Sequence | Tm* | Pool | Re-design strategy |
|---|---|---|---|---|---|---|---|
| 1_RIGHT | remove | 386..410 | minus | 5'-**CATCTTTAAGATGTTGACGTGCCTC**-3' | 66 | Pool1 | |
| 1_RIGHTv2 | add | 392..416 | minus | 5'-**AAGTGCCATCTTTAAGATGTTGACG**-3' | 65 | Pool1 | shifting position by 6-nt |
| 9_RIGHT | remove | 2883..2904 | minus | 5'-**ATGACAGCATCTGCCACAACAC**-3' | 68 | Pool1 | |
| 9_RIGHTv2 | add | 2886..2907 | minus | 5'-**TTTATGACAGCATCTGCCACAA**-3' | 64 | Pool1 | shifting position by 3-nt |
| 13_RIGHT | remove | 4143..4164 | minus | 5'-**ACCACAGCAGTTAAAACACCCT**-3' | 67 | Pool1 | |
| 13_RIGHTv2 | add | 4146..4167 | minus | 5'-**ATAACCACAGCAGTTAAAACAC**-3' | 60 | Pool1 | shifting position by 3-nt |
| 15_RIGHT | remove | 4675..4696 | minus | 5'-**AACAGAAACTGTAGCTGGCACT** -3' | 66 | Pool1 | |
| 15_RIGHTv2 | add | 4678..4699 | minus | 5'-**AGAAACAGAAACTGTAGCTGGC**-3' | 65 | Pool1 | shifting position by 3-nt |
| 21_LEFT | remove | 6168..6196 | plus | 5'-**TGGCTATTGATTATAAACACTACACACCC**-3' | 66 | Pool1 | |
| 21_LEFTv2 | add | 6165..6193 | plus | 5'-**TGGTGGCTATTGATTATAAACACTACACA**-3' | 65 | Pool1 | shifting position by 3-nt |
| 32_RIGHT | remove | 9835..9858 | minus | 5'-**AGCACATCACTACGCAACTTTAGA** -3' | 66 | Pool2 | |
| 32_RIGHTv2 | add | 9841..9864 | minus | 5'-**GGTAATAGCACATCACTACGCAAC** -3' | 65 | Pool2 | shifting position by 6-nt |
| 36_LEFT | remove | 10667..10688 | plus | 5'-**TTAGCTTGGTTGTACGCTGCTG**-3' | 68 | Pool2 | |
| 36_LEFTv2 | add | 10652..10680 | plus | 5'-**ATTACAGTTAATGTTTTAGCTTGGTTGTA**-3' | 62 | Pool2 | shifting position by 8-nt and extending 5' tail by 4-nt |
| 38_RIGHT | remove | 11669..11693 | minus | 5'-**CACCAAGAGTCAGTCTAAAGTAGCG**-3' | 67 | Pool2 | |
| 38_RIGHTv2 | add | 11675..11701 | minus | 5'-**ATCATAAACACCAAGAGTCAGTCTAAA**-3' | 63 | Pool2 | shifting position by 6-nt and extending the 5' end by 2-nt |
| 66_ LEFT | remove | 19845..19866 | plus | 5'-**GGGTGTGGACATTGCTGCTAAT**-3' | 68 | Pool2 | |
| 66_ LEFTv2 | add | 19845..19863 | plus | 5'-**GGGTGTGGACATTGCTGCT**-3' | 69 | Pool2 | removing 3-nt on the 3' end |
| 76_RIGHT | remove | 23193..23214 | minus | 5'-**ACACCTGTGCCTGTTAAACCAT**-3' | 67 | Pool2 | |
| 76_RIGHTv2 | add | 23241..23265 | minus | 5'-**TCTCTGCCAAATTGTTGGAAAGGCA**-3' | 69 | Pool2 | Primer3 |
| 89_LEFT | remove | 26836..26857 | plus | 5'-**GTACGCGTTCCATGTGGTCATT**-3' | 68 | Pool1 | |
| 89_LEFTv2 | add | 26833..26854 | plus | 5'-**CGCGTACGCGTTCCATGTGGTC**-3' | 74 | Pool1 | shifting position by 3-nt |
| 89_RIGHT | remove | 27203..27227 | minus | 5'-**ACCTGAAAGTCAACGAGATGAAACA**-3' | 66 | Pool1 | |
| 89_RIGHTv2 | add | 27209..27233 | minus | 5'-**ATAGTAACCTGAAAGTCAACGAGAT**-3' | 63 | Pool1 | shifting position by 6-nt |

* Tm for Q5 High-Fidelity DNA Polymerase calculated in NEB web site tool (https://tmcalculator.neb.com/)
